# Hampered motility promotes the evolution of wrinkly phenotype in *Bacillus subtilis*

**DOI:** 10.1101/288951

**Authors:** Anne Richter, Theresa Hölscher, Patrick Pausch, Tim Sehrt, Franziska Brockhaus, Gert Bange, Ákos T Kovács

## Abstract

**Background:** Selection for a certain trait in microbes depends on the genetic background of the strain and the selection pressure of the environmental conditions acting on the cells. In contrast to the sessile state in the biofilm, various bacterial cells employ flagellum-dependent motility under planktonic conditions suggesting that the two phenotypes are mutually exclusive. However, flagellum dependent motility facilitates the prompt establishment of floating biofilms on the air-medium interface, called pellicles. Previously, pellicles of *B. subtilis* were shown to be preferably established by motile cells, causing a reduced fitness of non-motile derivatives in the presence of the wild type strain.

**Results:** Here, we show that lack of active flagella promotes the evolution of matrix overproducers that can be distinguished by the characteristic wrinkled colony morphotype. The wrinkly phenotype is associated with amino acid substitutions in the master repressor of biofilm-related genes, SinR. By analyzing one of the mutations, we show that it alters the tetramerization and DNA binding properties of SinR, allowing an increased expression of the operon responsible for exopolysaccharide production. Finally, we demonstrate that the wrinkly phenotype is advantageous when cells lack flagella, but not in the wild type background.

**Conclusions:** Our experiments suggest that loss of function phenotypes could expose rapid evolutionary adaptation in bacterial biofilms that is otherwise not evident in the wild type strains.

## Background

Trait loss can have detrimental or beneficial consequences on the fitness of individuals. Eventually, loss in certain phenotypic attributes can have both negative and positive impact depending on the environmental conditions. However, impairment of certain paths might allow the evolution of new traits to compensate for the fitness loss. Such a trait loss and evolution can be easily detected in microbes that are able to promptly adapt to the selection pressure of their environment. Due to their rapid reproduction and pheno-or genotypic adaptation, evolution can be recognized even within few days. For example, *Pseudomonas fluorescens* rapidly adapts to static conditions and produces a microcosm at the air-medium interface, established by cellulose polymer overproducing derivatives [1]. These matrix overproducers, distinguished by their typical wrinkled colony morphotype in the laboratory can emerge in numerous bacterial species [2–4]. The evolution of these wrinkly morphotypes in *Pseudomonas* is governed by the altered bis-(3′-5′)-cyclic dimeric guanosine monophosphate (c-di-GMP) levels in the cells [5, 6]. It is suggested that the complexity and flexibility of the regulatory system around c-di-GMP facilitates adaptation to new environments [7].

Interestingly, elimination of the major c-di-GMP modulating components revealed several other mutational pathways allowing the appearance of wrinkly morphotypes. In addition, the appearance and fixation of newly evolved genotypes is facilitated by the spatial structure present in biofilms [8]. Various biofilm types are established by *Bacillus subtilis* under laboratory conditions, including pellicles at the air-medium interface [9–11]. *B. subtilis* cells inhabiting the biofilms are sessile and produce a matrix consisting of exopolysaccharides (EPS), protein fibers (TasA) and hydrophobin protein (BslA) [12–15]. Complex regulatory pathways ensure the mutually exclusive expressions of genes related to biofilm matrix production and motility [16, 17]. In addition to its role as the major repressor of biofilm formation, SinR also affects the expression of genes related to motility and cell separation collectively with other regulatory proteins [17]. Therefore, SinR has a central role in coordinating the exclusive expression of genes responsible for motile and sessile states.

While flagellum-dependent motility is not essential for the establishment of pellicles in *B. subtilis*, it facilitates the rapid formation of new biofilms [18]. Therefore, strains lacking motility are delayed in pellicle formation and are outcompeted by cells possessing the functional motility apparatus and exhibiting aerotaxis [18].

Rapid appearance of distinct *B. subtilis* morphotypes has been previously described in a 2-month-long batch culture experiment under static and shaken conditions [19]. Under both conditions, versatile morphotypes evolved including derivatives with reduced matrix production and linages with enhanced matrix expression. In the latter case, mutation in the *sinR* gene was identified by candidate-gene approach. Interestingly, mutations in *sinR* rapidly emerge in colonies of *B. subtilis* lacking SinI, an antagonist of SinR function [20]. Additionally, emergence of *sinR* mutants solves the problem of toxic galactose metabolites accumulation in the *galE* mutant, where elevated EPS production functions as a shunt for the toxic molecule [21]. Therefore, the adaptation pathway though *sinR* mutations appears to be a general denouement for numerous adaptation processes in *B. subtilis* biofilms.

Here, we study how the lack of functionally assembled flagella influences the evolution of wrinkly morphotypes in *B. subtilis* and demonstrate that matrix-overproduction caused by non-synonymous mutations in SinR primarily aids non-motile cells.

## Results and Discussion

### Evolution of wrinkly morphotypes in pellicles of *B. subtilis*

Lack of motility delays the establishment of *B. subtilis* pellicles [18]. During the previous study, when various biofilm competition experiments were performed using non-motile *B. subtilis* strains and colony-forming units (CFU) were assayed on LB plates, the appearance of a distinct colony phenotype was noticeable. The wrinkles and size of the observed colonies were clearly increased compared to their ancestors used for the study (Fig. 1a). Interestingly, these wrinkled spreader colonies (hereafter called WS morphotypes) were mostly apparent for the strain lacking the gene coding for the flagellin protein (i.e. *hag*). Therefore, a series of mutant strains used in our previous study (Hölscher *et al.*, 2015) was examined for the frequency of wrinkled derivatives during pellicle formation. Strains lacking various parts of the flagellum (flagellin (*hag*), hook (*flgE*), or basal body (*fliF*)), having disrupted regulation of motility (*sigD* mutant) or harboring a non-active flagellum (*motA* mutant) contained an increased amount of WS colonies during pellicle formation compared to the wild type (Fig. 1b).

**Fig. 1.**
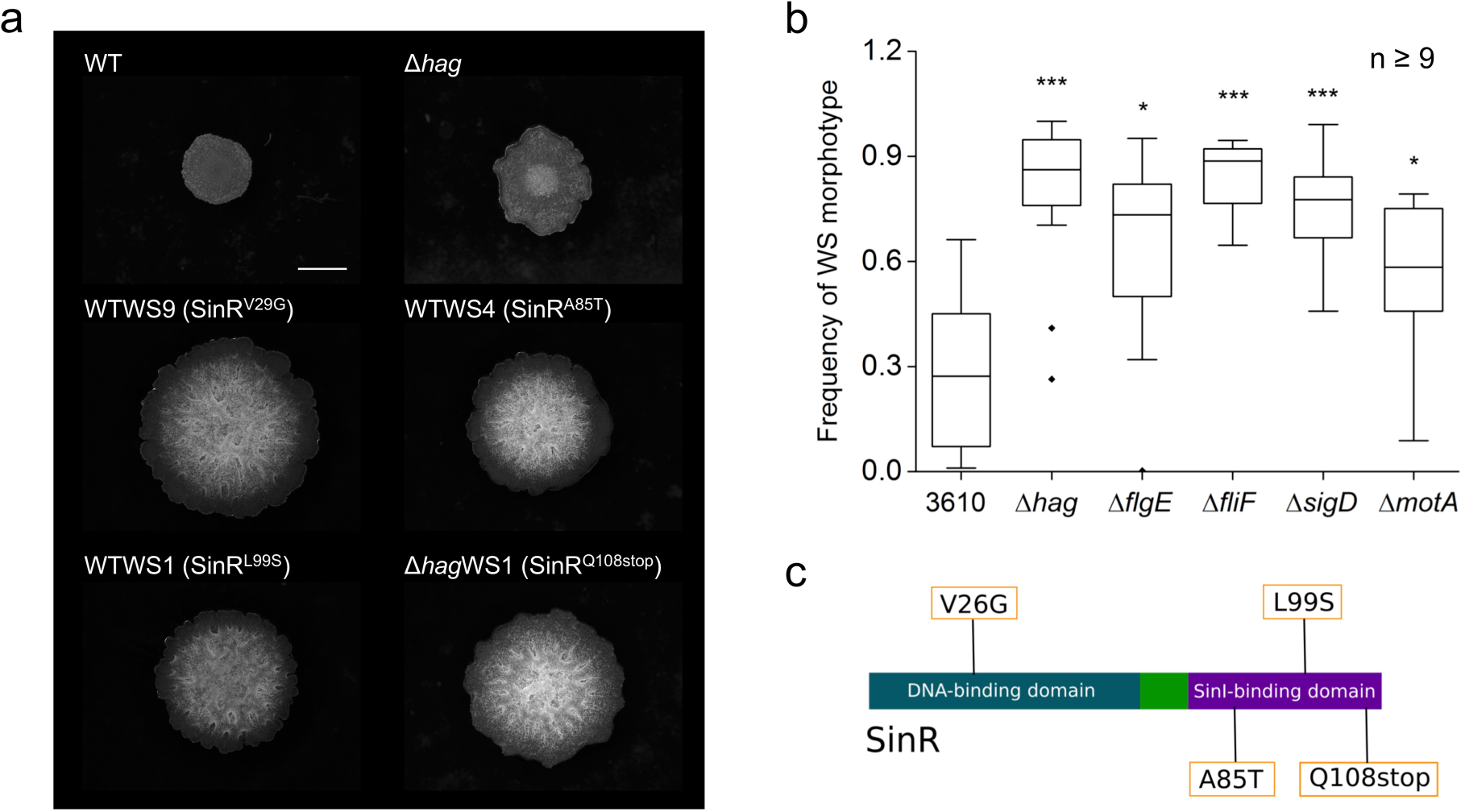
WS morphotypes exhibit elevated wrinkle formation and appear especially in non-motile mutant. (a) Microscopy images of WS morphotypes isolated from WT and Δ*hag* strains grown on LB medium. Representative images for the phenotype of each detected mutation, indicated in parentheses, are displayed. The scale bar represents 2 mm. (b) Relative frequency of WS morphotypes in the pellicle biofilms of various *B. subtilis* derivatives grown on MSgg medium. Y-axis shows the frequency of WS colony types (1 equals to 100% of the colonies being wrinkly). Boxes represent quartile 1-3, the line represents the median and whiskers indicate the upper and lower inner fence and dots represent outliers. Asterisks indicate significant differences (two sample Student’s t test assuming unequal variances: P values: Δ*hag,* 8.27•10-6; Δ*flgE*, 0.013; Δ*fliF*, 1.23•10-7; Δ*sigD*, 2.21•10-5, Δ*motA*, 0.0125; n ≥ 9). (c) Schematic representation of the SinR protein with domains and location of the detected mutations depicted.

### WS morphotypes harbor non-synonymous mutations in *sinR*

After isolation of ten WS morphotypes each from cultures of wild type and flagellin-lacking mutant (WTWS1-10 and Δ*hag*WS1-10, respectively), we analyzed the *sinR* gene encoding a major regulator of *B. subtilis* motility and biofilm formation, since wrinkle formation is among other factors associated with matrix production [12, 22]. SinR is a strong repressor of biofilm genes in *B. subtilis*, therefore gentle modification of such repressor function likely to impact the timing biofilm development. Sequencing revealed several non-synonymous substitutions resulting in SinR variants with the following changes in amino acid composition: V26→G, A85→T, L99→S and Q108→stop. V26G was located in the DNA-binding domain of SinR, whereas the other three mutations were found in the SinI-binding domain (Fig. 1c). All isolated morphotypes contained one of those mutations and exhibited a phenotype with increased wrinkles (Fig. 1a). Therefore, one of these mutations was probably sufficient to induce increased wrinkle formation in *B. subtilis*. In the absence of genome re-sequencing of the respective isolates, we cannot exclude the possibility that additional mutations present in the WS morphotypes that also contributes to the observed phenotypes. In our further experiments, we investigated strains representative for one of the detected mutations.

### WS morphotypes exhibit increased expression of matrix genes

As SinR is responsible for repression of the biofilm matrix genes, we examined their expression using strains containing the P*_tapA_*-*yfp* reporter, which is an indicator for the expression of the matrix operon *tapA-sipW-tasA*. Matrix gene expression of different WS morphotypes, a *sinR* mutant, as well as the ancestral strains of the wild type and *hag* mutant was qualitatively analyzed using fluorescence microscopy in LB medium that does not induce strong biofilms in *B. subtilis* (Fig. 2a). While P*_tapA_*-*yfp* expression was scarcely present in the wild type and hag mutant, it notably increased in the *sinR* mutant as well as in the WS morphotypes, whose cells occurred also primarily in chains. Mutation in *sinR* increases chain formation in planktonic cultures of *B. subtilis* as observed before [20]. Interestingly, the wild type and *hag* mutant showed detectable, although heterogeneous matrix gene expression in small clusters appearing after prolonged incubation, but they represented only a minor portion of the culture (Additional file 1: Figure S1). In contrast, the matrix gene expression of the *sinR* mutant and the WS morphotypes seemed to be homogeneously increased (Fig. 2a and Additional file 2: Figure S2). These results were confirmed by a quantitative analysis of the P*_tapA_*-*yfp* expression of the same strains over the course of 24 h (Fig. 2b). Here, the difference in expression level between *sinR* deletion mutant and the WS morphotypes became apparent, indicating that the mutations of the *sinR* variants did not abrogate the function of SinR completely. In addition, our measurements in shaken cultures suggest that the enhanced matrix expression in planktonic culture is likely contribute to the accelerated pellicle biofilm formation of the *hag* mutant.

**Fig. 2.**
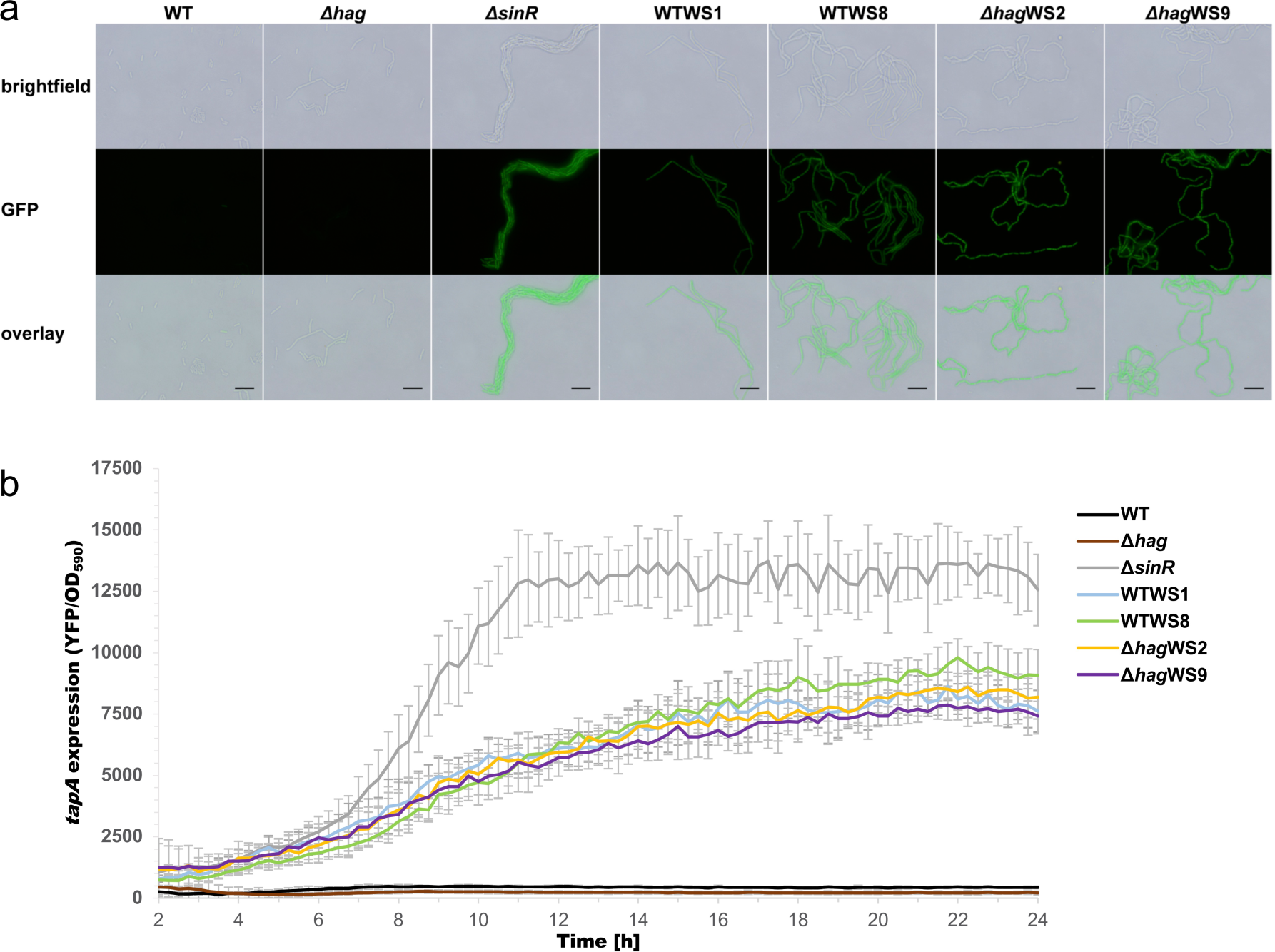
Increased matrix gene expression of WS morphotypes. (a) Representative confocal microscopy images of strains with P*tapA*-*yfp* reporter (false colored green) indicating matrix gene expression. Wild type, Δ*hag* and Δ*sinR* were compared to representative WS morphotypes of each wild type and Δ*hag* background harboring the mutations V26G (WTWS9 and Δ*hag*WS9) and L99S (WTWS1 and Δ*hag*WS2). The scale bar represents 10 µm. (b) OD normalized expression of P*tapA*-*yfp* of strains from (a) over time. Error bars represent the standard deviation.

### SinR-L99S differs from wildtype in its interaction with SinI and the SinR operator

Binding of the anti-repressor protein, SinI to SinR alters its specific DNA-binding properties. Due to the increased matrix gene expression, we hypothesized that the mutated SinR variants exhibit altered interaction properties with SinI, DNA or both. To test this hypothesis, the interaction of the SinR-L99S variant and SinI was investigated and compared to the interaction with the wild-type SinR. Variants SinR-V26G and SinR-A85T were not tested due to insolubility after overexpression under the conditions described in the Experimental Procedures section. To quantify the interaction between SinI and SinR or the SinR-L99S variant, we performed isothermal titration calorimetry (ITC), where SinR was titrated with SinI. In these experiments, a truncated version of SinI was used, a synthetic peptide consisting of amino acids 9-39 (encoded by *sinI*^9-39^). However, this short version was able to induce cell chaining when overexpressed in *B. subtilis* cells, similar to an overexpression of the full *sinI* (Additional file 3: Figure S3). Therefore, the short SinI version was sufficient to bind *in vivo* to SinR leading to a de-repression of the matrix genes and, thus formation of chains.

We observed tight binding of SinI to SinR with an apparent dissociation constant (K_D_) of approximately 7 nM and a stoichiometry of N = 1.2 +/- 0.02 assuming a one-site binding model (Fig. 3a). These data are in good agreement with the previously reported K_D_ of below 10 nM for the SinR/SinI interaction [23]. When we titrated the SinR L99S variant with the SinI peptide, we observed two distinct binding events (Fig. 3a). Applying a two-sites binding model, K_D_s of approximately 162 nM and 571 nM for the binding sites 1 and 2, respectively, were determined (Fig. 3a). These data suggest that binding of SinI to the SinR-L99S is weaker than for the wildtype and occurs in two different binding events. However, the summed stoichiometry of binding event 1 (N1 = 0.898 +/- 0.358) and 2 (N2 = 0.448 +/- 0.382) of N ≈ 1.35 suggests to us that no additional binding site is present. Interpreting our findings in the context of the SinI/SinR crystal structure delivers a plausible explanation for the bi-phasic binding of SinI to the SinR-L99S variant: In brief, SinR homo-dimerization with SinR and hetero-dimerization with SinI is mainly facilitated via the two C-terminal α-helices of SinR and is characterized by a hydrophobic core at the interface surrounded by polar interactions (Fig. 3c). Exchange of leucine to serine at position 99 at the border of the hydrophobic core creates a polar environment that should disturb the interaction with the unpolar interaction interface of SinI. This might also be the reason why we observed a two-phased binding event of the SinI peptide to SinR-L99S. It might well be that the SinI/SinR-L99S interaction occurs in a sequential manner and is first established by the N-terminal α-helix of SinI, before the C-terminal α-helix interlocks at the opposed site at the altered dimer interface. Closer inspection of the thermodynamic parameters revealed that SinI binding to wildtype SinR is entropy driven, as suggested by the positive ΔS of 24.7 cal mol^-1^ deg^-1^ and negative ΔH of −3752 +/- 135.6 cal mol^-1^ and might therefore be largely established by hydrophobic interactions. In contrast, binding of SinI to the SinR-L99S variant occurs in two steps with binding event 1 being characterized by a positive ΔS of 17.9 cal mol^-1^ deg^-1^ and a negative ΔH of −3928 +/- 1450 cal mol^-1^ indicative of an interaction that is mainly established via hydrophobic interaction. However, binding event 2 is characterized by a negative ΔS of −22.9 cal mol^-1^ deg^-1^ and a negative ΔH of −1.524E4 +/- 1.24E4 cal mol^-1^. Hence, the mechanism of binding event 2 seems enthalpic and entropic, and might involve expulsion of structured H_2_O molecules from the binding site. This observation supports the model in which SinI association to SinR-L99S occurs in two steps with the second binding step being affected by an impaired hydrophilic interface caused by the polar serine in position 99.

**Fig. 3.**
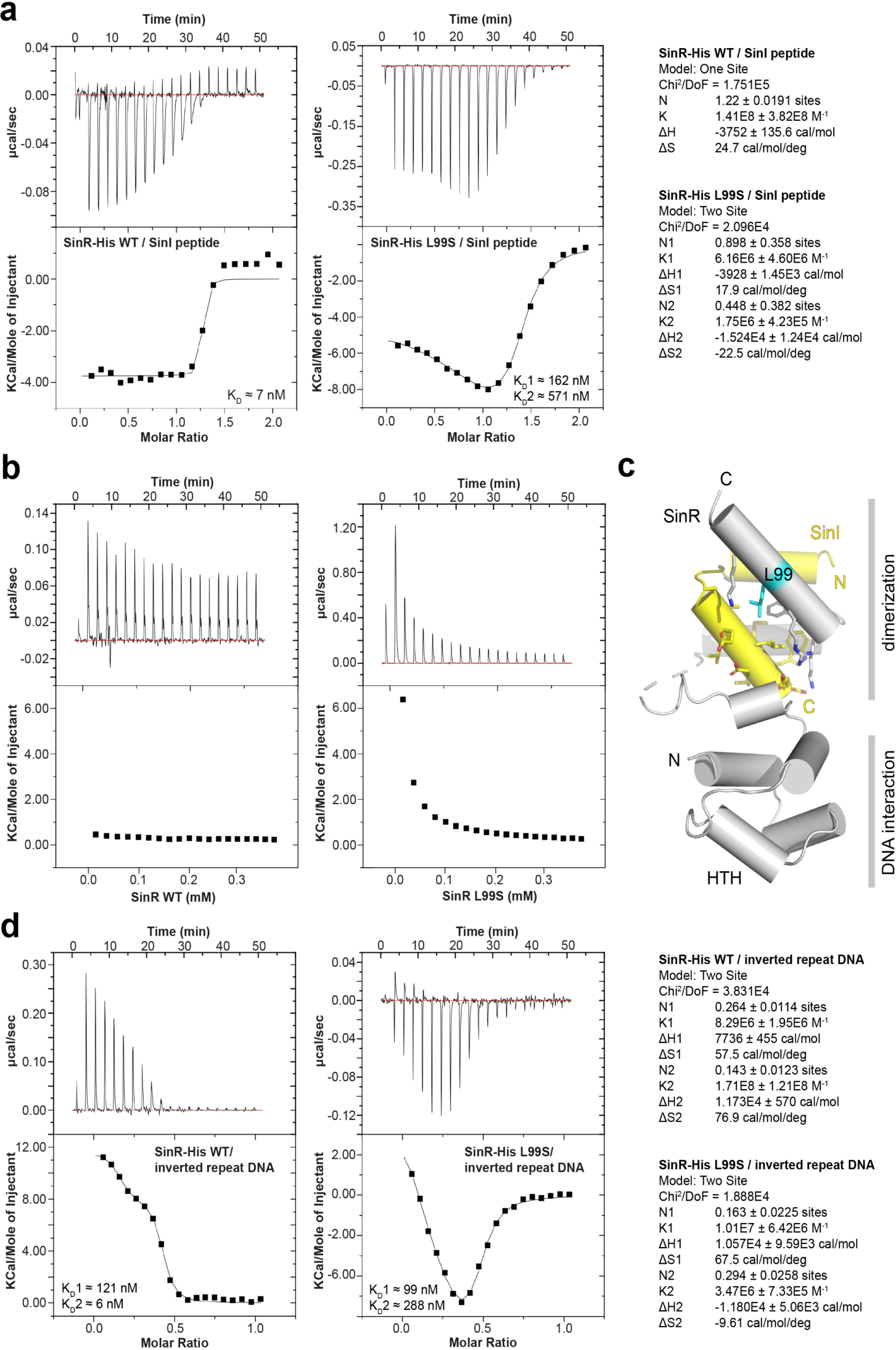
SinR-L99S differs in its interaction with SinI and DNA. (a) ITC measurments of the interaction between the SinI peptide and SinR (left thermogram) and SinR-L99S (right thermogram). Derived thermodynamic parameters are shown on the right site. (b) ITC complex dissociation experiments of SinR (left thermogram) and SinR-L99S (right thermogram). (c) Cartoon representation of the *B. subtilis* SinR/SinI complex crystal structure (PDB-ID: 1B0N; Newman et al. 2013). SinR is colored in grey and SinI is colored in yellow. Leucine 99 (cyan) and the sourrunding SinR/SinI interface region is shown in stick representation. N and C indicate N-termini and C-termini, respectively. (d) ITC measurments of the interaction between the inverted repeat DNA and SinR (left thermogram) and SinR-L99S (right thermogram). Derived thermodynamic parameters are shown on the right site.

Next, we reasoned that the presence of serine at position 99 might affect the formation and stability of the SinR tetramer. We therefore analyzed the oligomeric properties of SinR and SinR-L99S by analytical size exclusion chromatography (SEC), revealing that wildtype SinR and SinR-L99S both occur almost exclusively as tetramers (Additional file 4: Figure S4a). However, the L99S mutation might affect tetramer stability. Therefore, we assayed the tetramer dissociation properties of the wildtype and SinR-L99S by ITC. To do so, wild-type or mutant SinR protein was titrated into buffer. Strikingly, while the wild-type SinR did not dissociate, the L99S variant showed reproducibly detectable reduction in tetramer stability upon rapid dilution (Fig. 3b). It might well be that this subtle defect has significant consequences at the functional level, i.e. the interaction of SinR with the DNA operator sequence. However, analytical SEC of reconstituted SinR/IR-DNA and SinR-L99S/IR-DNA complexes revealed that interaction of SinR or SinR-L99S with the IR-DNA SinR operator occurs in the tetrameric state (Additional file 4: Figure S4b). Taken together, these findings agree well with the binding model proposed by Newman and co-workers [23].

### SinR-L99S shows impaired interaction with the SinR operator

Having demonstrated that SinR-L99S interacts as tetramers with the IR-DNA SinR operator, we wondered whether the interaction of SinR-L99S with IR-DNA differs from wildtype SinR in respect to the binding affinity. We performed ITC on SinR, respectively SinR-L99S, in the sample cell and titrated the IR-DNA into the cell. ITC revealed that interaction of SinR with the IR-DNA occurs in a bi-phasic manner. We therefore applied a two-sites binding model to the data, revealing that the two binding events are characterized by K_D_s of approximately 121 nM and 6 nM for the binding sites 1 and 2, respectively (Fig. 3d). Our study confirms previous experiments stating the high-affinity interaction of SinR with IR-DNA [23]. The summed stoichiometry of binding event 1 (N1 = 0.264 +/- 0.0114) and 2 (N2 = 0.143 +/- 0.0123) of N ≈ 0.41 further suggests to us that one IR-DNA fragment is able to interact with two SinR proteins, which is in good agreement with the crystal structure of SinR in complex with the IR-DNA SinR operator (Additional file 5: Figure S5) [23]. As shown above by analytical size exclusion chromatography, interaction of the IR-DNA occurs at the tetramer, likely in a 4:2 stoichiometry (SinR:IR-DNA). It is therefore unclear if the two observed binding events represent cooperative binding events at the same IR-DNA fragment or cooperative binding events at the opposed sites of the tetramer, relayed via the dimerization domain of SinR.

Strikingly, ITC titration of the L99S variant SinR protein with the IR-DNA fragment revealed that the interaction was impaired, as suggested by the K_D_s of approximately 99 nM for site 1 and 288 nM for site 2 (Fig. 3d). Interestingly, while binding event 1 is comparable to the wildtype interaction for binding site 1 in the affinity and thermodynamic parameters ΔS1 and ΔH1, the thermodynamic parameters for binding event 2 changed from a ΔS2 of 76.9 cal mol^-1^ deg^-1^ and a ΔH2 of 1.173E4 +/- 570 cal mol^-1^ for the wildtype interaction to a ΔS2 of −9.61 cal mol^-1^ deg^-1^ and a ΔH2 of −1.18E4 +/- 5.06E3 cal mol^-1^ for the SinR-L99S/SinI interaction. In summary, the interaction of SinR with the IR-DNA is changed by serine in position 99 at the SinR dimerization interface in that it seems to alter not only the affinity for the IR-DNA at the DNA binding site in the second binding step, but also the mechanism by which the interaction is established. Hence, the interaction of wild-type SinR with the IR-DNA is characterized by positive cooperativity, while the SinR-L99S/IR-DNA interaction is impaired and displays features of negative cooperativity. This might result in de-repression of SinR target operons in case of the SinR-L99S variant and is in good agreement with the observed phenotype.

### WS morphotype in Δ*hag* background is advantageous during pellicle establishment

Next, we were interested in whether the WS morphotypes success during colonization of the air-liquid interface is altered, since they appeared frequently under these conditions. To test this, we competed the WS morphotype SinR^L99S^ (WTWS1 and Δ*hag*WS2) against their respective ancestral strains (i.e. wild type or *hag* mutant) under conditions allowing pellicle formation and detected the strains using constitutively expressed fluorescence reporters. Figure 4a shows that both wild type and WS morphotype were equally successful in colonization of the air-liquid interface, which was comparable to the controls with competitions of the same strain. In contrast, the competition between the *hag* mutant and its derived WS morphotype revealed that the Δ*hag*WS morphotype was able to outcompete the *hag* mutant during pellicle establishment (Fig. 4a). Interestingly, in each competition with the WS morphotype, the structure of the pellicle displayed a higher spatial segregation of cells than the wild type control competition that was visible as patches of red or green fluorescent regions. In the Δ*hag* background, this effect can be explained by the inability to mix due to lack of flagella (comparable with the Δ*hag* control competition as described previously by [18]). However, in the WTWS morphotypes, this assortment could be due to a reduced motility that accompanies the increased matrix production. Indeed, *sinR* mutant has been previously described to display diminished motility [24]. Additionally, the high amount of produced matrix might add to the adhesiveness of the cells, so that they clump together after cell division. The semi-quantitative analysis of the signal abundance for these competition experiments confirmed the superior surface colonization of the WS morphotypes compared to the ancestor in the Δ*hag* but not the wild type background (Fig. 4b). In spite of the observed comparable fitness of WT and WTWS, *de novo* evolution of WTWS is still noteworthy (see Fig. 1b), although clearly lower than in the *hag* mutant background. In addition, the Δ*hag*WS morphotype was able to establish a thin pellicle at the air-liquid interface faster than the *hag* mutant when compared as single strain cultures (Additional file 6: Video S1).

**Fig. 4.**
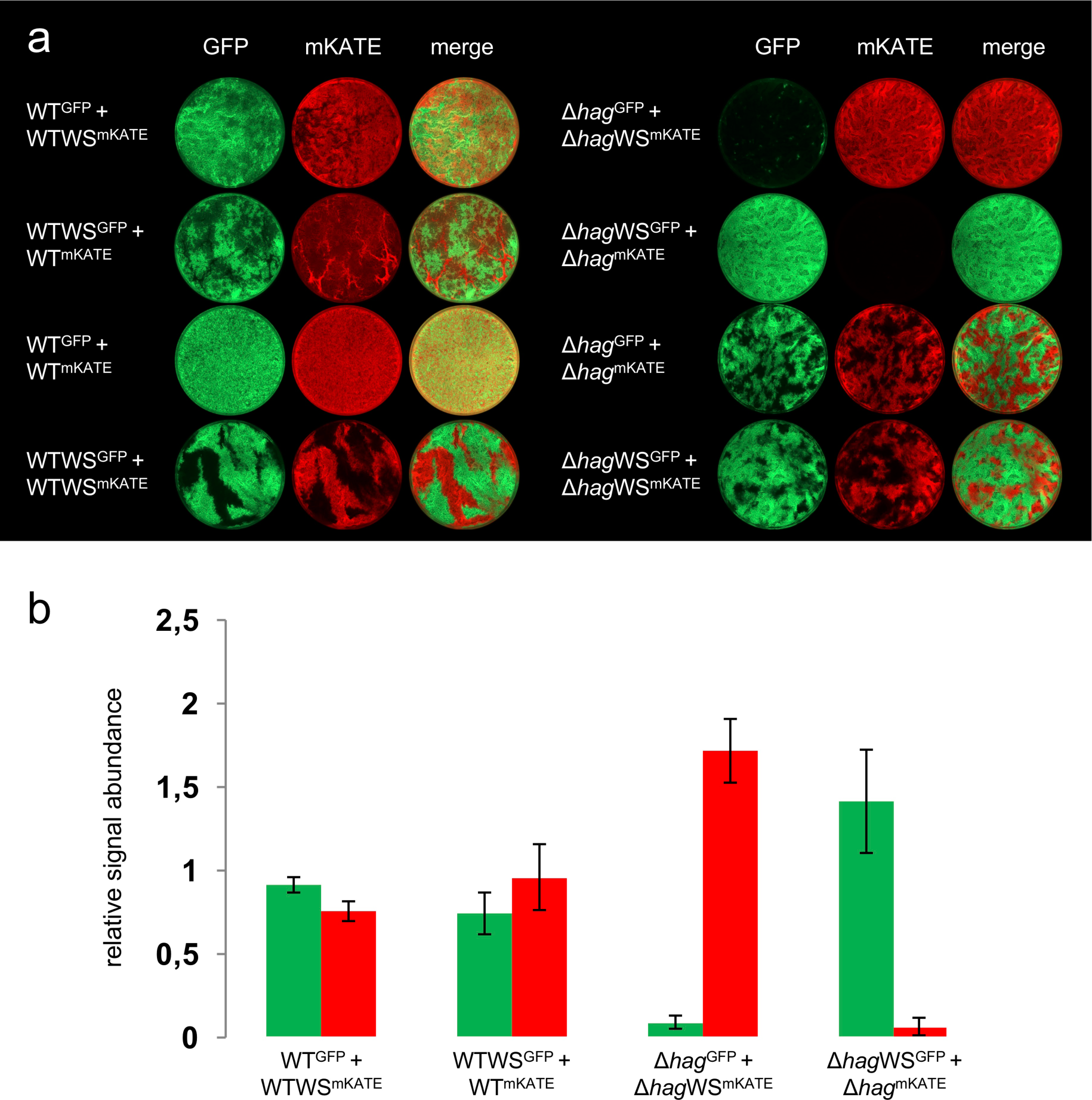
WS morphotypes outcompete ancestor in Δ*hag* but not wild type background. (a) Microscopy images of competitions between green (GFP) and red (mKATE) fluorescently labelled wild type (left) or *hag* mutant (right) and derived WS morphotypes (SinRL99S). False colored images of wells of a 24-well plate (diameter: 16 mm) are displayed. Control competitions with swapped fluorescent reporters (2nd row) and between otherwise identical strains (3rd and 4th row) were performed for each. (b) Semi-quantitative analysis of relative signal abundance for competitions of ancestor and WS morphotype from (**a**) (two sample Student’s t test assuming unequal variances: WTGFP + WTWSmKATE P = 0.005; WTWSGFP + WTmKATE P = 0.134; Δ*hag*GFP + Δ*hag*WSmKATE P = 2.8•10-4; Δ*hag*WSGFP + Δ*hag*mKATE P = 0.0027; n = 4). Error bars represent the standard deviation.

### Selection pressure, not mutation rate is responsible for WS morphotype appearance

To investigate if an increased mutability of the *hag* mutant compared to the wild type is responsible for the primary occurrence of WS morphotypes in this strain, we determined the frequency of streptomycin-resistant mutants in both strains. Since the frequency of mutants in wild type and Δ*hag* with respective mean values of 8,05⋅10^−6^ (standard deviation: 5,01⋅10^−6^) and 8,26⋅10^−6^ (standard deviation: 2,35⋅10^−6^) were comparable, we could exclude the mutation rate as reason for the frequent appearance of WS morphotypes. Because of the advantage of the WS morphotype in surface colonization in the Δ*hag* background, we conclude that there the selection pressure was high enough to result in mutations aiding the establishment of a pellicle at the air-liquid interface. This was probably caused by decreasing oxygen levels towards the bottom of the vessel as well as the limited number of cells that reach the liquid surface due to the lack of swimming motility, which is present in the wild type. Besides or because of an increased adhesiveness, the elevated matrix production of the WS morphotypes possibly results in a higher buoyancy, which counterbalances the lack of swimming and facilitates the fast surface colonization as indicated by Video S1 in Additional file 6. Therefore, an increased selection pressure was likely responsible for the primary occurrence of WS morphotypes in the *hag* mutant. Although suppressor mutants of *sinR* were found under different conditions in the laboratory [19, 21], in nature, the observed mutations in SinR probably appear less frequent since most environmental isolates of *B. subtilis* are motile.

## Conclusions

Bacteria possess the ability to adapt to a huge variety of environments and conditions, with often impressive solutions to their challenges. To overcome their disadvantage of slow surface colonization in small numbers during pellicle establishment, non-motile *B. subtilis* strains develop suppressor mutations in *sinR*, encoding an important regulator and part of the intricate regulatory network governing biofilm formation in *B. subtilis*. These mutations alter its DNA- and protein-binding properties, leading to increased production of the biofilm matrix. In turn, matrix overproduction allows a faster surface colonization than the ancestor, successfully outcompeting it. Therefore, *B. subtilis* provides an interesting example of bacterial adaptability and resourcefulness in conditions with a specific selective pressure as well as it might explain their success on earth.

## Methods

### Media composition and culturing conditions

All the strains used in this study are listed in Additional file 7: Table S1. For cloning, mutant generation and colony morphology experiments, strains were cultivated in Lysogeny Broth medium (LB-Lennox, Carl Roth, Germany; 10 g l^-1^ tryptone, 5 g l^-1^ yeast extract and 5 g l^-1^ NaCl) supplemented with 1.5 % Bacto agar if required. To select for wrinkly phenotypes in pellicles, strains were pre-grown in LB medium overnight and diluted 1:100 in MSgg medium (5 mM potassium phosphates buffer (pH 7), 100mM MOPS, 2 mM MgCl_2_, 700 µM CaCl_2_, 100 µM MnCl_2_, 50 µM FeCl_3_, 1 µM ZnCl_2_, 2 µM thiamine, 0.5 % glycerol, 0.5 % glutamate [9]). When 2 ml culture was incubated in a 24-well plate under static conditions at 30 °C, pellicles were formed at the air-medium interface after 72 h. If appropriate, the following antibiotics were used: Kanamycin (Km, 5 µg ml^-1^), Lincomycin + Erythromycin (MLS, 12.5 µg ml^-1^ + 1 µg ml^-1^, respectively), Chloramphenicol (Cm, 5 µg ml^-1^), Spectinomycin (Spec, 100 µg ml^-1^) and Ampicillin (Amp, 100 µg ml^-1^).

### Isolation of WS strains and genetic analysis of the *sinIR* locus

Wild type or various mutant strains of *B. subtilis* were grown in MSgg medium under static conditions and adequate dilutions were spread on LB agar plates to obtain single colonies. Colonies with wrinkly phenotypes were counted. Selected colonies were cultivated in LB medium, genomic DNA was extracted using EURex Bacterial & Yeast Genomic DNA Kit (Roboklon GmbH, Berlin, Germany), the *sinIR* locus was PCR amplified using primers oTB98 and oTB99 (see Additional file 8: Table S2 for oligonucleotide sequences), and PCR products were sequenced (GATC GmbH, Cologne, Germany).

### Constructions of plasmids and strains

*B. subtilis* strains (using DK1042 based strains that is naturally competent version of NCIB3610 [25]) were obtained via natural competence transformation using genomic or plasmid DNA [26]. Strains with constitutively expressing green-or red-fluorescent reporters were obtained by transforming genomic DNA from 168hyGFP or 168hymKATE, respectively [27]. Biofilm specific reporter strains were created using genomic DNA from DL821 harboring a P*_tapA_*-*yfp* reporter construct [28]. To overexpress the *sinI* and *sinI*^9-39^ genes, the full and the truncated genes were obtained using oligonucleotides oTB124-oTB125 and oTB126-oTB127 (Additional file 8: Table S2), respectively, digested with *Hin*dIII and *Sph*I enzymes, and cloned into the corresponding sites of pDR111 (kind gift from David Rudner), resulting in pTB695 and pTB696, respectively. The obtained plasmids were verified using sequencing and introduced into *B. subtilis* DK1042 using natural competence [26]. Transformants were selected on LB plates with appropriate antibiotics. When appropriate, successful transformation was validated using the fluorescence reporter activity of the strains or amylase-negative phenotype on 1 % starch agar plates.

To overexpress *sinR*, the wild type gene was amplified by PCR from *B. subtilis* 168 genomic DNA (sequence identical in NCIB 3610) using SinR_NcoI_F and SinR_H6_BamHI_R primers (Additional file 8: Table S2), harboring a C-terminal hexa-histidine tag. The fragment was digested with *Nco*I and *Bam*HI restriction enzymes and cloned into a pET24d vector for overexpression in *E. coli*. Mutagenesis of *sinR* was performed in a two-step PCR mutagenesis with the respective mutagenesis primer pairs (Additional file 8: Table S2) and subsequent cloning as described above.

### Microscopy analysis of competition experiments and *sinI* overexpression

For competition experiments, fluorescently labeled strains were used. The fluorescent-based semi-quantitative measurement was previously shown to correlate with colony forming unit-based detection [18]. The pre-grown GFP and mKATE labeled strains were mixed at equal optical density and diluted 1:100 in MSgg medium. After 72 h of growth, the fluorescence intensity was measured using an infinite F200PRO plate reader (TECAN Group Ltd, Männedorf, Switzerland). Bright field, green- and red-fluorescence images of the pellicles were taken with an Axio Zoom V16 stereomicroscope (Carl Zeiss, Jena, Germany) at 3.5x magnification equipped with a Zeiss CL 9000 LED light source, HE eGFP filter set (excitation at 470/40 nm and emission at 525/50 nm), HE mRFP filter set (excitation at 572/25 nm and emission at 629/62 nm), and an AxioCam MRm monochrome camera (Carl Zeiss, Jena, Germany). The exposure times were set to 0.01 s, 1 s and 3 s for bright field, green- and red-fluorescence, respectively. ImageJ (National Institute of Health, Bethesda, MD, USA) was used for background subtraction and channel merging.

For *sinI* overexpression, strains TB697 or TB698 were pre-grown in LB medium overnight, diluted 1:100 in fresh LB medium, and incubated in the absence or presence of 0.1 mM IPTG for 4 h. Samples were added to microscopy slides containing a thin layer of 1 % agarose, glass coverslips were placed on the samples and the cells were visualized using MOTIC BA310E phase contrast microscope equipped with a 100x/1.25 PHASE EC-H objective and a MOTICAM 3 camera (VWR, Darmstadt, Germany).

### Reporter assays

To monitor biofilm coupled gene expression, wild type and selected wrinkly isolates harboring the P*_tapA_*-*yfp* reporter construct were pre-grown on LB agar plates. One colony was inoculated in 3 ml liquid LB medium and incubated for 5 h at 37 °C and 225 rpm, diluted to DO_600_ of 0.1 in LB medium, and 200 μl aliquots of the cultures were inoculated into a 96-well plate. The samples were incubated for 24 h at 30 °C with continuous shaking among measurements, and optical density and fluorescence was recorded every 15 minutes.

For single-cell fluorescence microscopy, one colony from overnight grown plate was inoculated in 3 ml liquid LB broth and incubated for 5 h at 37 °C and 225 rpm. 5 μl of culture was spotted on a microscope slide coated with 0.8 % agarose, covered with a cover slip and examined under the fluorescence microscope (Olympus Bx51; 100× oil objective). Images were captured using bright light (exposure time 15 ms) and fluorescence light using the GFP filter (exposure time 500 ms).

### Fluctuation assay

To determine the mutation rate, a fluctuation assay was performed with wild type and *hag* mutant as described in [29], except for the use of MSgg medium (n = 48).

### Protein expression and purification

Constructs of pET24sinR^WT^ and their mutagenized variants were transformed into *E. coli* BL21(DE3) for protein overexpression. Proteins were overexpressed in 1 l LB autoinduction media (1.8% w/v lactose) shaking at 30 °C overnight. Cells were harvested the next morning and resuspended in 20 ml buffer A (20 mM HEPES/NaOH pH 8.0, 500 mM NaCl, 40 mM imidazole). Cells were lysed two times with a M-110L Microfluidizer (Microfluidics) and centrifuged at 20,000 r.p.m. for 20 min at 4 °C to remove cell debris. The supernatant was applied onto a 1 ml HisTrap HP column (GE Healthcare) for Ni-NTA affinity chromatography. The column was washed with 15 column volumes of buffer A and proteins were eluted with 5 ml buffer B (20 mM HEPES/NaOH, pH 8.0, 500 mM NaCl, 500 mM imidazole). Proteins were concentrated to 1 ml and further purified by size-exclusion chromatography using a HiLoad 26/60 Superdex 200 gel-filtration column in buffer C (20 mM HEPES/NaOH, 500 mM NaCl).

### Isothermal titration calorimetry

ITC experiments were performed on a MicroCal ITC 200 instrument (GE Healthcare). The SinI^9-39^ peptide (SinI protein from amino acid 9 to 39) was synthesized with a free amine at the N-terminus and free acid group at the C-terminus (peptides&elephants GmbH, Potsdam, Germany). The peptide was dissolved in an appropriate volume of buffer C, which was used for the purification of SinR^WT^ and SinR^L99S^. Concentrations were determined by measuring the A_280_ using a NanoDrop Lite spectrophotometer (Thermo Scientific). For the ITC SinR/SinI interaction experiment, 200 μl of SinR^WT^ or SinR^L99S^ (25 μM) was added to the sample cell and 250 μM SinI^9-39^ peptide solution were titrated in at 25 °C for a total of 20 injections, each separated by 150 s, consisting of 0.2 μl SinI^9-39^ peptide for the initial injection and 2 μl for the following 19 injections. For the SinR dissociation ITC experiment, 200 μl of buffer C was added to the sample cell and 2 mM SinR^WT^ or SinR^L99S^ were titrated in at 25 °C for a total of 20 injections, each separated by 150 s, consisting of 0.2 μl SinI^9-39^ peptide for the initial injection and 2 μl for the following 19 injections. For the ITC SinR-inverted repeat DNA interaction experiment, 200 μl of SinR^WT^ or SinR^L99S^ (30 μM) was added to the sample cell and 150 μM inverted repeat DNA was titrated in at 25 °C for a total of 20 injections, each separated by 150 s, consisting of 0.2 μl SinI^9-39^ peptide for the initial injection and 2 μl for the following 19 injections. The inverted repeat primers were prior to the experiment dissolved in buffer C and annealed by heating to 95 °C for 5 min and a subsequent controlled cooling down to 10 °C for 1 h, using a PCR cycler. ITC data were processed using the Origin ITC software (OriginLab) and thermodynamic parameters were obtained by fitting the data to a one set of sites binding model or a two sets of sites binding model, depending on the data.

## Declarations

### Availability of data and materials

All bacterial strains are available from the corresponding author.

#### Authors’ Contributions

ÁTK design of the study; AR, TH, TS, FB, PP, ÁTK conducted experiments; AR, PP, GB, ÁTK interpreted the data; TH and ÁTK wrote the manuscript; AR, PP, GB corrected the manuscript

## Ethics approval and consent to participate

Not applicable.

## Consent for publication

Not applicable.

---

**Figure. S1.**
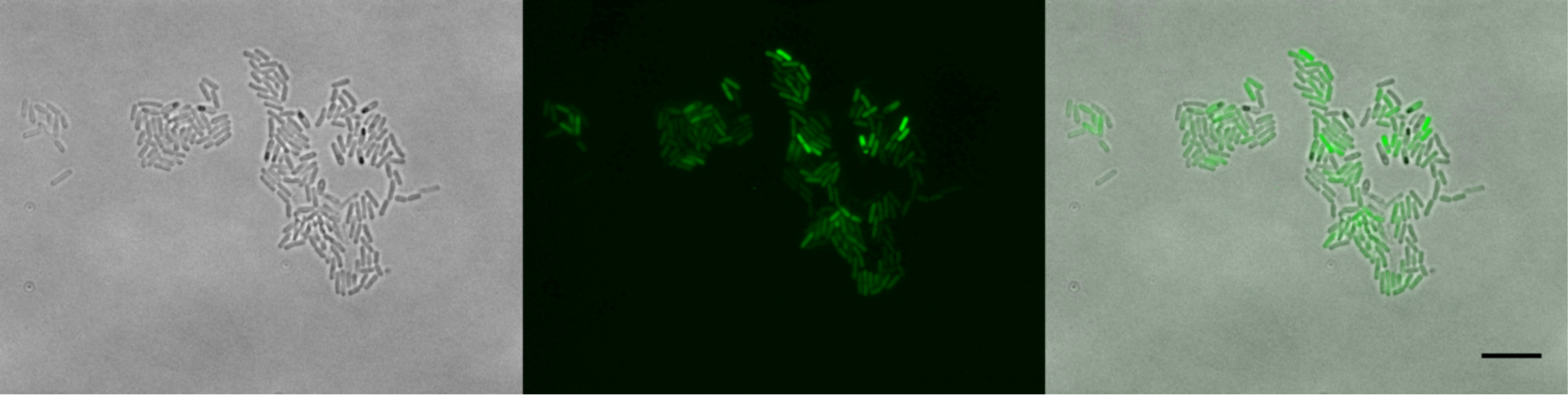
Prolonged incubation of wild-type *B. subtilis* in LB medium results in aggregation with increased, but heterogeneous *tapA* expression. Representative microscopy images of strains harboring the P*^tapA^-yfp* reporter in wild type background. Images were recorded as in Fig. 2, but after longer (>12 h) incubation in LB medium. The presented aggregates were observed in addition to homogeneously dispersed cells similar to those observed in Fig. 2. The scale bar represents 10 µm.

**Figure. S2.**
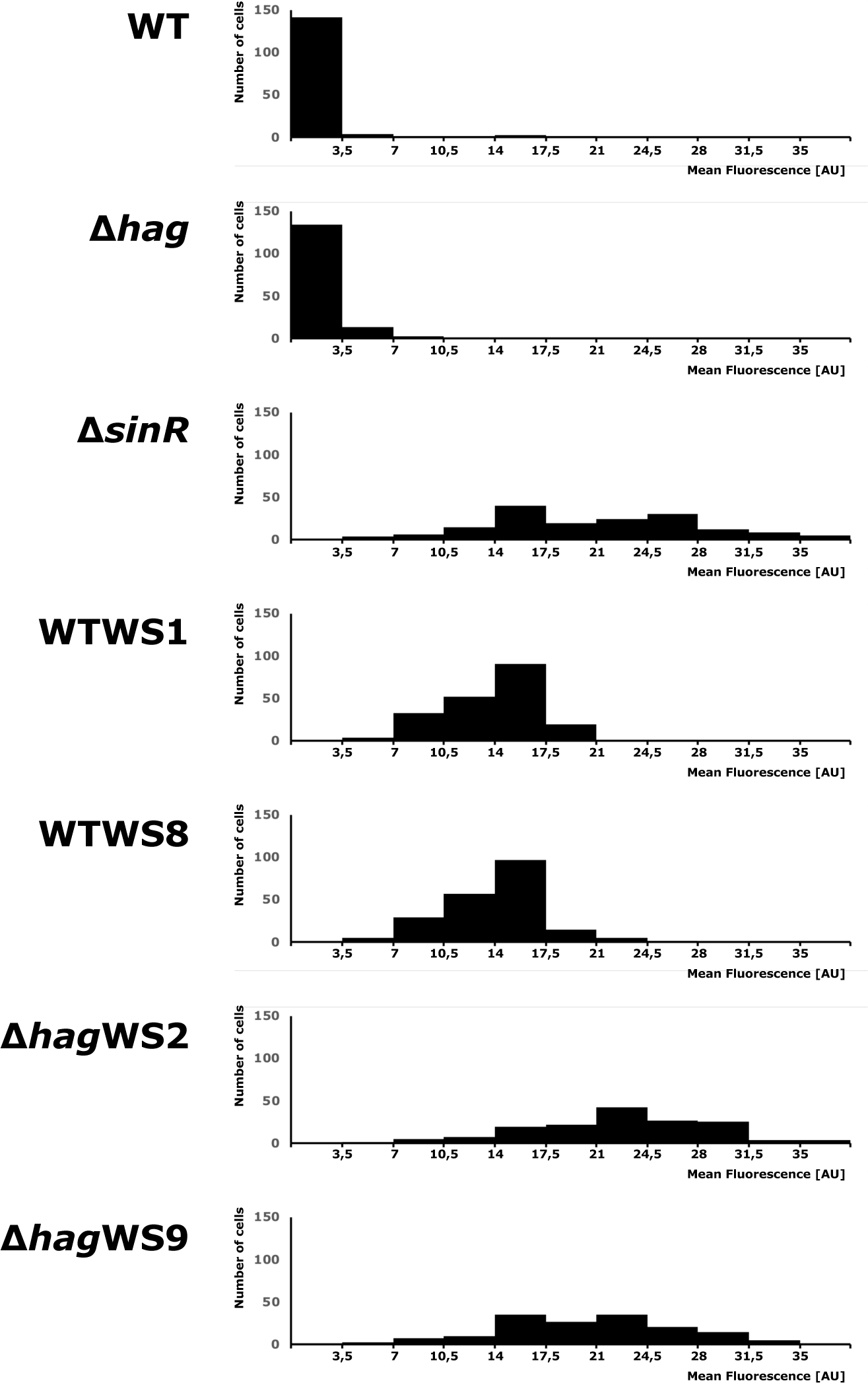
Expression from the P*_tapA_*-*yfp* reporter detected at single cell level. Histograms were created based on images similar to Fig. 2a after randomly selecting 150 bacterial cells and detecting mean fluorescence using Image J (version 2.0.0-rc-68/1.52e). Background fluorescence was determined for each image based on 5 random selected positions where no cells were visible, and the average value was subtracted from each fluorescence data measured for the given image. X axis indicates mean fluorescence (arbitrary units), while Y axis denotes the number of cells.

**Figure. S3.**
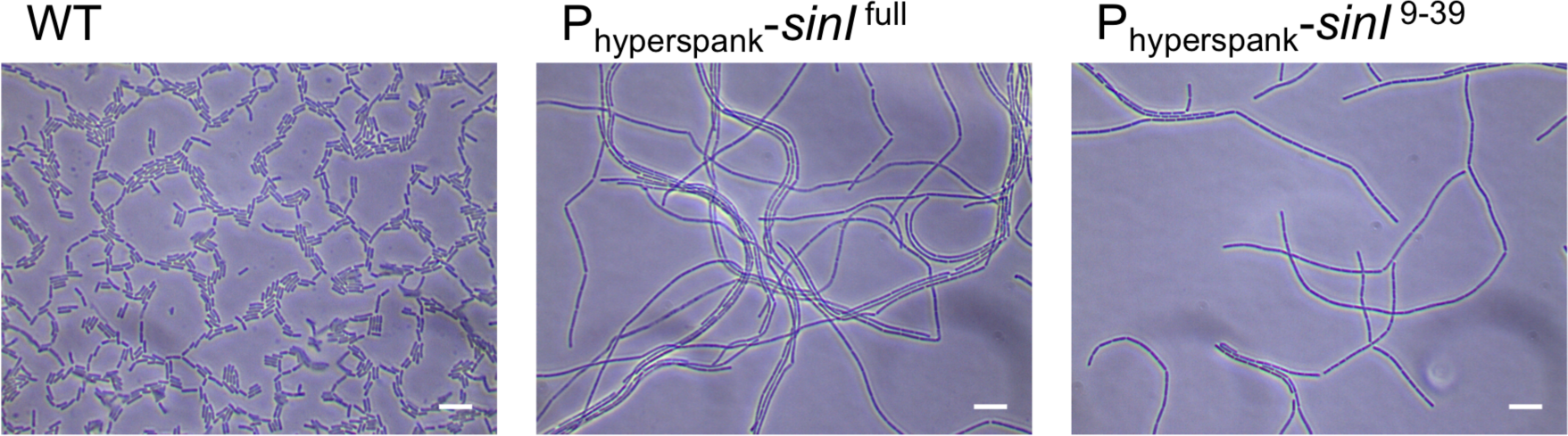
Truncated SinI supports cell chaining. Microscopy images of wild type and two *B. subtilis* strains harboring a *sinI* overexpression construct of the full gene (*sinI*^full^) or a truncated version (*sinI*^9-39^). The scale bar represents 10 µm.

**Figure. S4.**
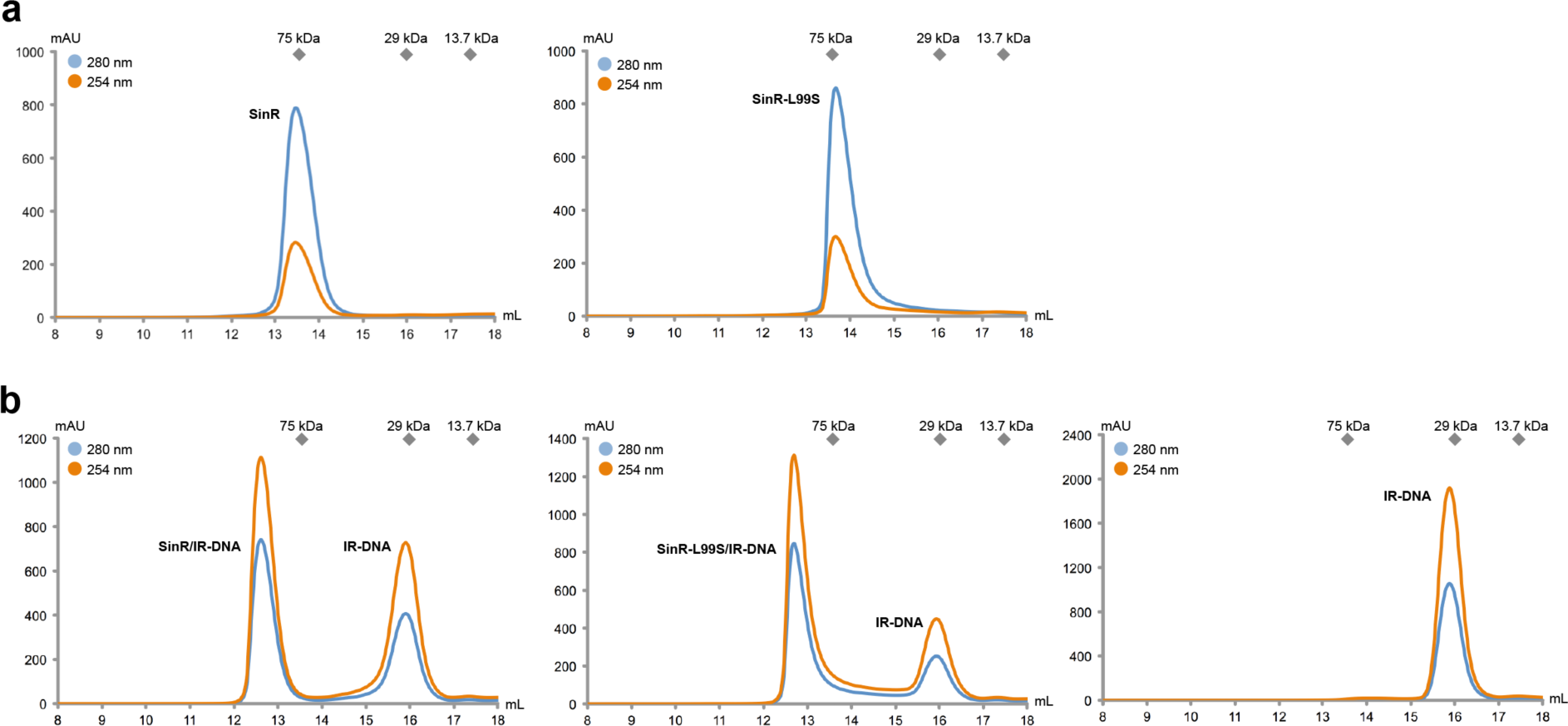
SinR and SinR-L99S function as tetramers. (a) Analytical size exclusion chromatograms of SinR (left) and SinR-L99S (right). Runs were performed on a Superdex 200 Increase 10/300 GL column. The absorbance was recorded at 254 nm (red curve) and 280 nm (blue curve) in mAU (arbitrary units). (**b**) Analytical size exclusion chromatograms of the reconstituted SinR/IR-DNA complex (left), the SinR-L99S/IR-DNA complex (middle) and the individual IR-DNA duplex. Runs were performed on a Superdex 200 Increase 10/300 GL column. The absorbance was recorded at 254 nm (red curve) and 280 nm (blue curve) in mAU (arbitrary units).

**Figure. S5.**
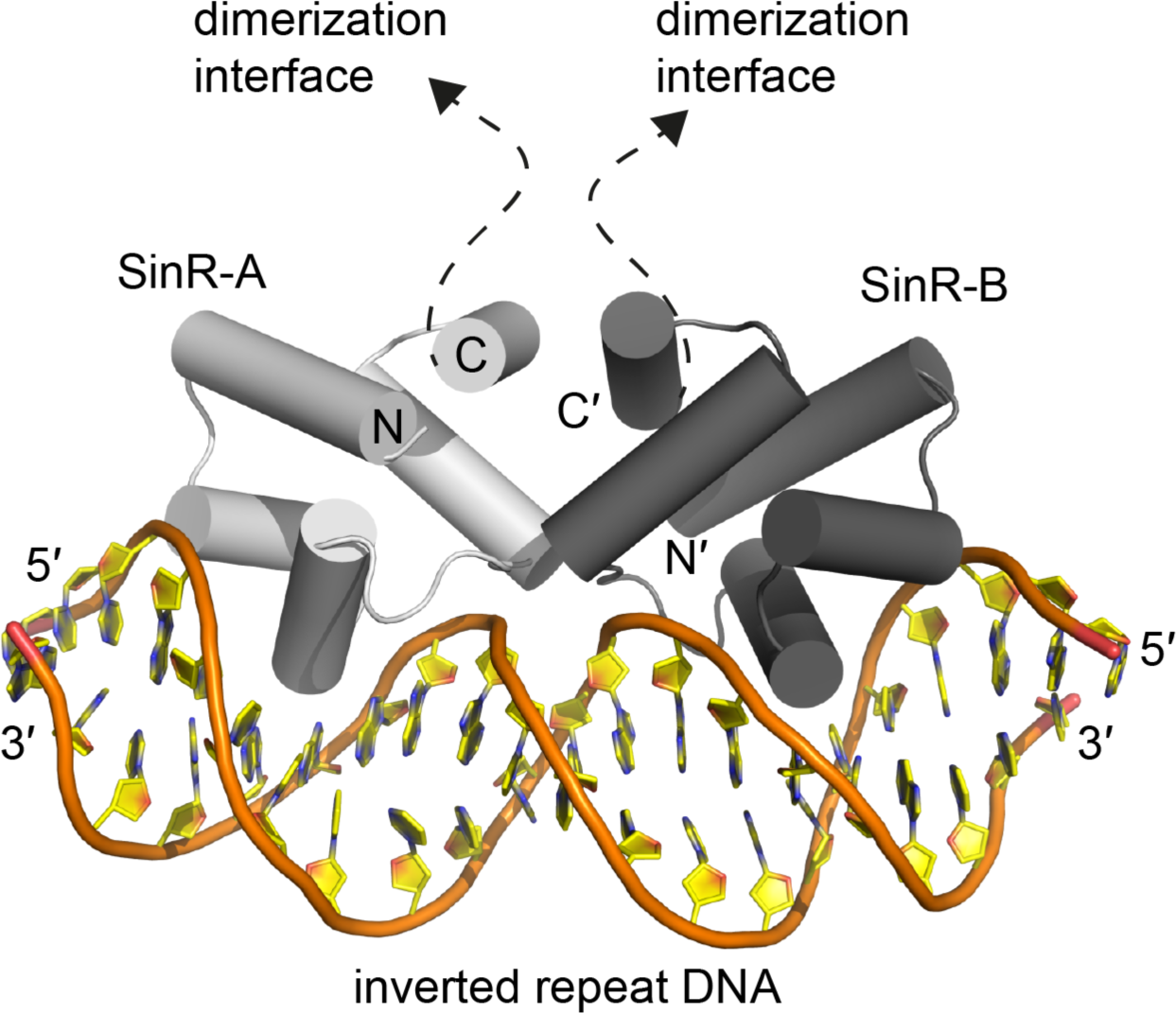
Crystal structure of the SinR/DNA complex. The crystal structure of the *B. subtilis* SinR/IR-DNA complex is shown in cartoon representation (PDB-ID: 3ZKC; [23]). Two SinR proteins (SinR-A, colored in light grey and SinR-B, colored in dark grey) bind via their N-terminal DNA interaction domain to inverted repeat DNA. N and C indicate N-termini and C-termini, respectively.

Video S1. Timing of pellicle initiation in *B. subtilis* Δ*hag* and its WS derivative differs. Pellicle formation of *B. subtilis* Δ*hag* (mixture of TB36 (Δ*hag*^GFP^) and TB37 (Δ*hag*^mKATE^), left petri-dish) and its WS derivative (mixture of TB775 (Δ*hag*WS2^GFP^) and TB776 (Δ*hag*WS2^mKATE^), right petri-dish) is shown in MSgg medium. Cultures were grown in 35 mm diameter Falcon petri dishes at 30 °C and images were recorded every 30 min.

**Table S1.**
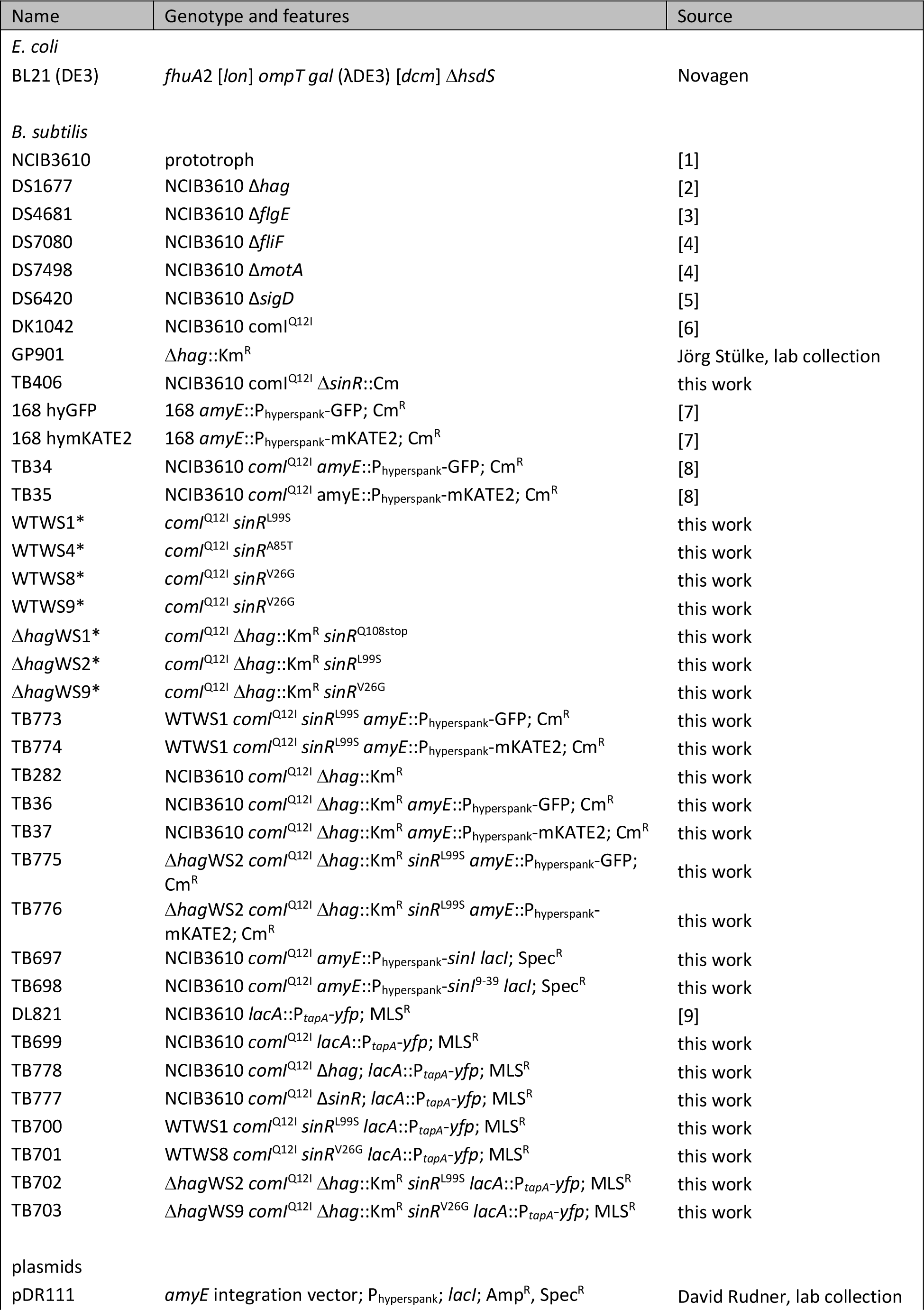
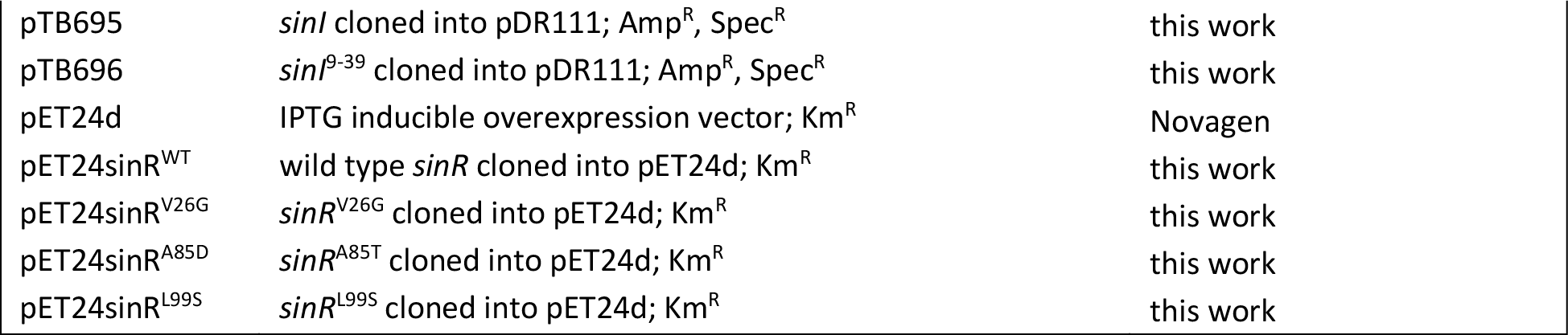
Strains and plasmids used in the current study. Strains labeled with *might contain additional mutations

**Table S2.**
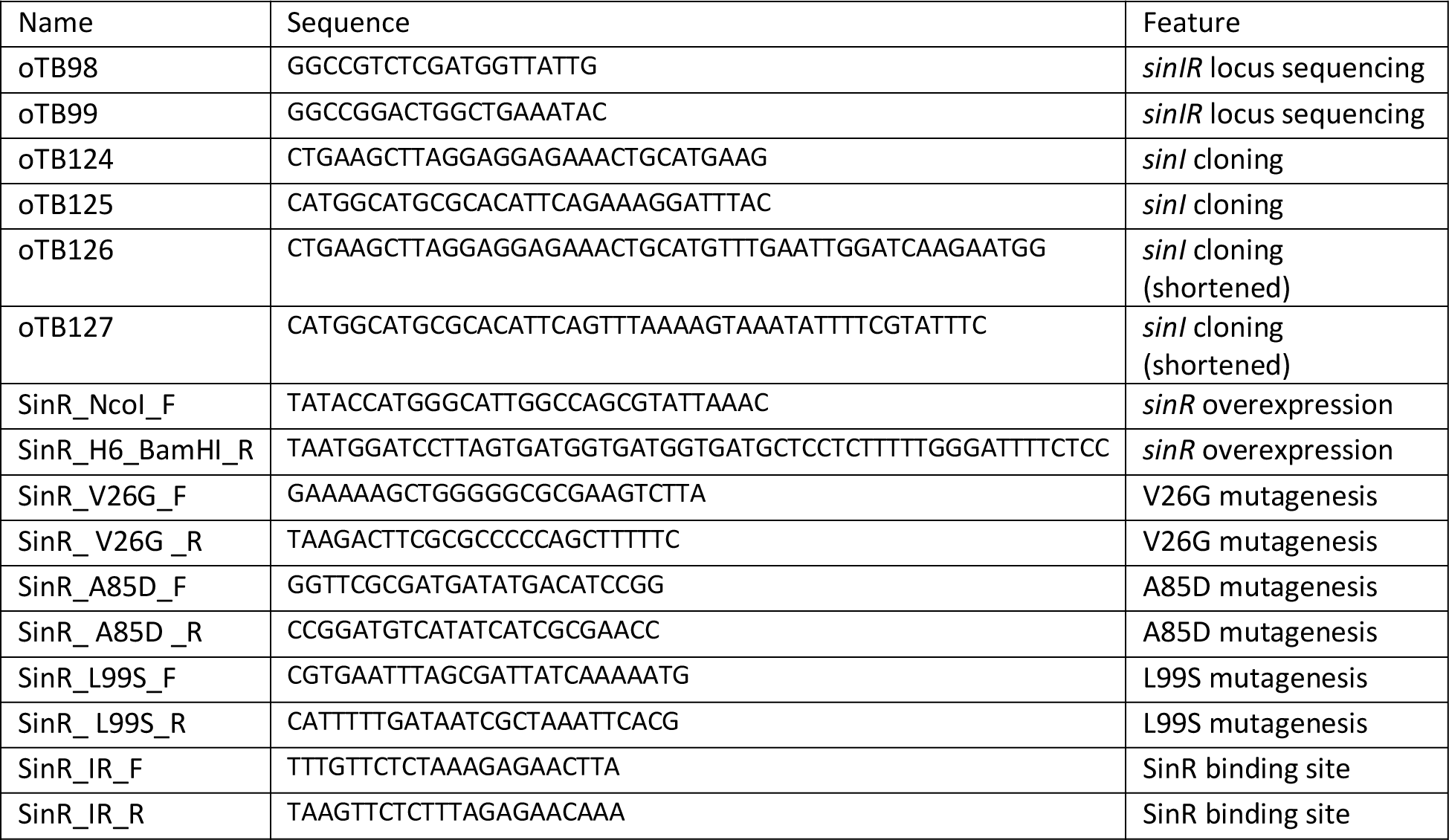
Oligonucleotides used in the current study

